# Coarse-grained Force Fields from the Perspective of Statistical Mechanics: Better Understanding the Origins of a MARTINI Hangover

**DOI:** 10.1101/2020.06.25.171363

**Authors:** Zack Jarin, James Newhouse, Gregory A. Voth

## Abstract

The popular MARTINI coarse-grained model is used as a test case to analyze the adherence of top-down coarse-grained molecular dynamics models (i.e., models primarily parameterized to match experimental results) to the known features of statistical mechanics for the underlying all-atom representations. Specifically, the temperature dependence of various pair distribution functions, and hence their underlying potentials of mean force via the reversible work theorem, are compared between MARTINI 2.0, Dry MARTINI, and all-atom simulations mapped onto equivalent coarse-grained sites for certain lipid bilayers. It is found that the MARTINI models do not completely capture the lipid structure seen in atomistic simulations as projected onto the coarse-grained mappings, and that issues of accuracy and temperature transferability arise due to an incorrect enthalpy-entropy decomposition of these potentials of mean force. The potential of mean force for the association of two amphipathic helices in a lipid bilayer is also calculated and, especially at shorter ranges, the MARTINI and all-atom projection results differ substantially. The former is much less repulsive and hence will lead to a higher probability of MARTINI helix association in the MARTINI bilayer than occurs in the actual all-atom case. Additionally, the bilayer height fluctuation spectra are calculated for the MARTINI model and – compared to the all-atom results – it is found that the magnitude of thermally averaged amplitudes at intermediate length scales is quite different, pointing to a number of possible consequences for realistic modeling of membrane processes. Taken as a whole, the results presented here can point the way for future coarse-grained model parameterization efforts that might bring top-down coarse-grained models into better agreement with the statistical mechanics of the actual all-atom systems they aspire to represent.

## Introduction

There are two general approaches to parameterizing a coarse-grained (CG) model: bottom-up and top-down.^1-4^ The bottom-up approach implies a direct correspondence to a finer resolution (e.g., an all-atom) model that corresponds to the system of interest. In contrast, the top-down approach has a direct connection mainly (or only) to the experimental system of interest. In the bottom-up case, the parameterization scheme results in a state-dependent CG potential. On the other hand, the top-down potential is dependent on the data fitted and the scheme used. Rigorous studies of bottom-up methods have identified two major issues in their CG parametrization: Transferability and representability. Generally, the transferability problem relates to the application of a CG model to conditions away from the original parameterization conditions, and it limits their extensibility to different thermodynamic state points.^5-6^ Separately, representability issues are rooted in key mathematical differences in the CG versus all-atom expressions.^7^ Perhaps the most common example is that bottom-up CG models may incorrectly capture the pressure, isothermal compressibility, and/or certain other observables due to a difficulty in transitioning the definitions of observables between the all-atom and CG representations.^8^ On the other hand, these problems have largely not been investigated or ignored in top-down CG models, where it is usually assumed (arguably wrongly) that the standard expressions for observables can simply be used in the CG model and that the CG model is somehow transferable (e.g., through a broader parameter fitting scheme).

MARTINI models are commonly used top-down CG models, most notably for their computational efficiency. The significant speedup compared to all-atom (AA) models arises, in part, from the reduced resolution, i.e., mapping approximately four heavy atoms to one CG site or “bead”. Specifically, the reduced representation significantly decreases the computational effort needed for a given simulation by reducing the force calculations per time step, while also increasing the diffusion coefficients of the system by a factor of ∼ 3-6, further increasing the temporal and spatial sampling of a system.^9-10^ Additionally, by employing an integration time step significantly longer than typical atomistic systems further increases the efficiency while maintaining a degree of numerical accuracy when sampling the underlying energy landscape. Despite the possible drawbacks from a loss of rigor, the overall computational saving inherent in MARTINI simulations has proven attractive when attempting to model complex biomolecular systems, which require longer length and time scales than are readily available with fully atomistic models.

Additionally, the MARTINI parameter set necessary to run the diverse set of molecules is reduced to only the polar (P), nonpolar (N), apolar (C), and charged (Q) CG bead types, which leads to a simpler parameterization and optimization strategy. The original parameters were fit using the octanol-water partition coefficient to capture the free energy of transfer from the hydrophobic region of a lipid bilayer to the fully hydrated exterior.^10^ The simplicity of bead types and potentials gives MARTINI models a modular and additive representation that provides an initial guess for CG force field parameters for new molecules. This ease of use – without the user needing to invest much further thought and which might reduce the time to publication – seems to be another factor for the widespread use of the MARTINI model for complex biomolecular and related systems.

Although MARTINI is widely utilized, there are possible drawbacks in accuracy due to lack of a rigorous CG mapping from the atomistic level, and the consequent top-down parameterization. The approximation of 4 heavy atoms-to-1 CG bead can be a poor description for chemically similar lipids because lipids with different tails are represented by the same MARTINI models, e.g., a single model represents C12:0 dilauroyl (DLPC) and C14:0 dimyristoyl (DMPC) tails. There is clearly no rigorous correspondence between a MARTINI model and the atomistic model in this case. The MARTINI model is, therefore, a general model with looser phenomenological connections to the underlying atomistic models. This is further reinforced in the parameterization scheme, which is based on matching experimental data and, for the most part, not the atomistic interactions and observables.

The challenge in mapping 4 heavy atoms-to-1 CG bead is most apparent when describing water, i.e., 4 waters-to-1 CG bead. As has been discussed elsewhere,^11-14^ MARTINI water has several problems, including freezing point, diffusion, and hydration behavior. This is apparent in analyses of lipid bilayer structure but is not the focus of the current study. Instead, we focus here more on the issues of transferability and representability as they pertain to certain known statistical mechanical properties of lipid bilayers. We also analyze the lipid bilayer undulation spectrum of the MARTINI model and compare it to analyses from CG mapped atomistic trajectories.

In this study, we use the frequently studied C18:1 dioleoyl (DOPC) plus cholesterol bilayers as test cases because lipid bilayers provide adequate configurational sampling at both MARTINI and atomistic resolutions, enabling clear comparisons to fundamental statistical mechanical quantities, e.g., radial distribution functions (RDFs). The MARTINI simulation of lipid bilayers is also thought to be the least fraught with difficulty because as opposed, e.g., to CG protein simulations the lipid bilayers are driven by “simpler” amphipathic assembly forces which should, in principle, be well-captured by a top-down CG model parameterized (in essence) on such forces. We also present a more applied example of differences between the MARTINI and atomistic resolutions: the lateral association of two amphipathic helices embedded in a lipid bilayer. We characterize the lateral helix association free energy by quantifying the potential of mean force (PMF) with respect to the center of mass (CoM) distance between two H0 helices of endophilin. Lastly, there are some properties of lipid bilayers calculable from molecular and CG simulations that do have a direct experimental analogue (i.e., bending modulus from the fluctuation spectra). These quantities are described in the Theory section and presented in Results.

## Theory

Generally speaking, CG potentials are parameterized to capture the behavior of a system using a reduced representation compared to the underlying finer-grained (e.g., atomistic) system. Since the reduced number of degrees of freedom in the CG system cannot fully capture the entropy of the system, a state dependence arises, and the CG potential is conditionally parameterized for a specific ensemble. The state dependence becomes an issue when a CG potential is used outside of the specific ensemble for which the CG potential is necessarily parametrized (e.g., at a higher temperature, with different composition, or at an increased surface tension).

In the case of the MARTINI CG force field, one might expect the same transferability problem to arise considering the approximate 4 to 1 mapping scheme and the interaction parameterization scheme detailed elsewhere.^10^ Here, we look first at temperature dependence. More specifically, we calculate the enthalpic and entropic contributions to a given PMF, *W*(*R*), between two CG sites spaced apart by a distance *R*, as calculated from radial distribution function, *g*(*R*), via the reversible work theorem such that^15^ *W*(*R*) = − *k*_*B*_*T* ln *g*(*R*).^5-6, 16^ The PMF is in fact defined as a conditional free energy, so it must be decomposable into the two contributions We will calculate a midplane-projection of the pairwise PMF in a single leaflet and then its decomposition into the enthalpic and entropic contributions (see Methods for further computational details).^16^ This decomposition of the PMF gives direct insight into the nature of the temperature dependence and provides a deeper understanding of the missing or incorrect entropic and enthalpic contributions to the MARTINI CG interactions compared with the underlying mapped atomistic scale results. This statistical mechanical insight into the nature of the MARTINI CG interactions can provide a better understanding of how issues of physical accuracy and transferability may appear (and can potentially be dealt with) in top-down CG models.

Representability issues in CG models, often more opaque, are related to the roles of resolution and the parameterization scheme in defining observables and calculating them.^6-8^ We seek here to highlight the nature of representability as it applies to a particular lipid observable, the height fluctuation spectrum. The spectrum is calculated through a discrete Fourier transform of the midplane lipid bilayer shown below:^17-18^ 

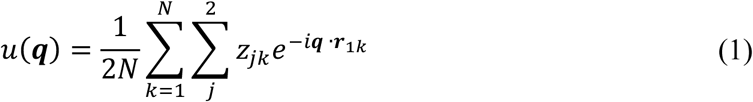

where *u* is the Fourier coefficient, ***q*** is the two dimensional reciprocal space vector, *N* is the number of lipids, *z*_*j,k*_ is the z-position of a *k*th lipid in *j*th leaflet and *r*_*j,k*_ is the (*x,y*) position of the *k*th lipid in the *j*th leaflet. The height fluctuation spectrum from the molecular dynamics simulation shown in Equation (1) can be related to height fluctuation spectrum from common Canham-Helfrich-type continuum models, shown below^19-20^ 

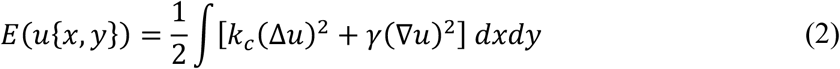

where *E* is the free energy, *k*_*c*_ is the bending modulus, and *γ* is the area compressibility. After a continuous Fourier transform and the decoupling of harmonic Fourier modes through equipartition, the ensemble average of the Fourier modes can be related to the bending modulus such that 

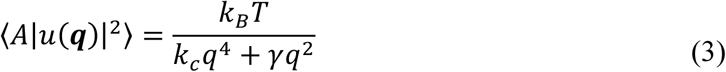

where *A* is the instantaneous area of the bilayer patch, *k*_*B*_ is Boltzmann’s constant, and *T* is temperature. This technique is widely used to relate molecular simulations to an analytical, continuum representation of an elastic sheet.^10, 17-18^ At zero surface tension, the height fluctuation spectrum provides a direct connection to continuum theory, and a means to estimate the bending modulus using one parameter fits to the low wavelength coefficients of the height fluctuation spectrum.^17^ This method has been applied to all-atom and CG models alike using the same formula, with the general expectation that the spectrum converges to the continuum result at low wavenumbers (i.e., long wavelength modes in molecular simulations behave similarly to continuum models). However, the relationship between the resolution of the atomistic and CG models and the height fluctuation spectrum at shorter wavelengths is poorly understood.

We thus expect CG mapped all-atom (i.e., all-atom models mapped to the resolution of MARTINI models) and MARTINI fluctuation spectra to converge to Helfrich-like behavior in the low wave number regime, where the corresponding length scale is much larger than a single phosphorus atom (or phosphate bead). However, we are interested in the intermediate lengthscale regimes, which can be related to molecular level detail as the corresponding length scale is of the order of these simulations. The height fluctuations in the intermediate regime, 0.8 nm^−1^ to 6 nm^−1^, are of specific interest because they are on the length scale of membrane-mediated interactions and have implications for using MARTINI to understand association of lipids with proteins, as well as the influence of the motions of the former on the conformations of the latter, when embedded in the membrane.

## Methods

### A. Simulation Details

All simulations were performed using GROMACS molecular dynamics (MD) simulation suite (versions 5.1.4 and 2016.3)^21^ using the CHARMM36 atomistic force field,^22^ MARTINI 2.0, and Dry MARTINI CG force fields.^10, 23^ The simulations for structural analysis of DOPC and cholesterol were 270 DOPC molecules and 68 cholesterol molecules with ∼36 waters per lipid (12,180 water molecules). The initial structures were generated using the CHARMM-GUI.^24-26^ Each simulation was run for a microsecond at 1 atm using the Nose-Hoover thermostat with the coupling of 1ps and the semi-isotropic Parrinello-Rahman barostat^27^ at 1.0 atm with the coupling of 5ps. The simulations to study the membrane undulation spectrum had 1152 DOPC molecules for the smaller 20nm by 20nm box size and 28,800 DOPC molecules for the larger 100nm by 100nm box size. The amphipathic helix simulations contain two H0 helices of endophilin in a bilayer of 140 DOPC lipids hydrated by 4,200 water molecules and 0.15 M of KCl. After embedding, umbrella sampling was used to restrain the center of mass (CoM) distances every 0.1 nm from 1.5nm to 3nm using a harmonic umbrella potential with force constant 15 kJ/mol implemented in Plumed v2.3.^28-29^ Finally, weighted histogram analysis method was used to reproduce the PMF from the final 100ns of a 150ns run at each window.^30-31^

### B. xy-projection of Per Leaflet Radial Distribution Function

We analyzed the *xy*-projection of the RDF for lipids of each leaflet as a way to understand the lateral association of lipids and cholesterol in each leaflet. First, the *xy*-projection was used because lipid bilayers are not spherically symmetric and the normalization of a radial distribution function assuming spherical symmetry results in an RDF that decays to 0 rather than the typical 1. Second, distinguishing between leaflets provides deeper insights into the near 0 distance behavior. The *xy*-projection for both leaflets shows significant nonzero behavior at distances near 0 corresponding to the lipid in the opposite leaflet. The contribution from lipid in the opposite leaflet is not significant, but it complicates distinguishing overlap between a choline-group of a lipid and a hydroxyl-group of cholesterol in the same leaflet, which is significant. Thus, the per leaflet *xy*-projection provides insight into the lateral behavior of the bilayer and correctly attributes *xy*-overlap to headgroup interaction instead of effects due to the opposite leaflet. Given the significant amount of cholesterol flip-flop, we assigned each lipid to a leaflet by determining if the tail CG bead (C4A bead of DOPC or C2 bead of cholesterol) is at least 0.5 nm above or below the head group bead (NC3 bead of DOPC or ROH bead of cholesterol). A Comparison between leaflets also provides a means of determining convergent behavior as both leaflets of a symmetric bilayer should show similar behavior. Notably, beads that are not interacting in the same leaflet can share *xy*-position. Thus, their corresponding RDF will be relatively flat. Similarly, little information can be gleaned from the RDF of a head group bead (e.g., choline bead) and corresponding tail bead (e.g., terminal hydrophobic bead). The RDF remains near 1, as the position of the head group bead has a small effect on the corresponding position of the tail bead in disordered lipid bilayers.

Next, we use the reversible work theorem to determine the corresponding PMF from the RDF. This PMF is a measure of the stability of two lipid CG beads sharing the same *xy*-position in a leaflet of the bilayer. For lipid beads with the same preferred *z*-position in the membrane (i.e., overlapping projected *z*-density), the PMF is a proxy for the lateral association energy of the two beads. After assuming negligible heat capacity change, the PMFs can be decomposed across different temperatures according to: 

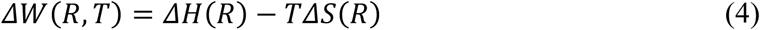

Numerically, we evaluate the decomposition by performing a linear fit of the PMF and temperature at a given distance value *R* to determine the slope (i.e., the entropy term) and *y*-intercept (i.e., the enthalpic term). The enthalpic and entropic terms can subsequently be plotted at each distance value and compared across models to gain a deeper understanding of the lipid association behavior and bead-wise interactions in the bilayer. The results were calculated using both the MARTINI CG models and the all-atom MD with the atomic coordinates mapped on the CG sites via a CoM mapping.^32-33^

## Results

### A. Lateral Association

When considering the lateral organization of cholesterol in a bilayer and the main driving forces, we first analyzed the previously described per leaflet *xy*-projection of the RDF. A subset of the RDFs shown in Fig. 1 using the MARTINI bead naming convention (i.e., NC3 corresponds to the choline group of the headgroup).^10^

**Figure 1:**
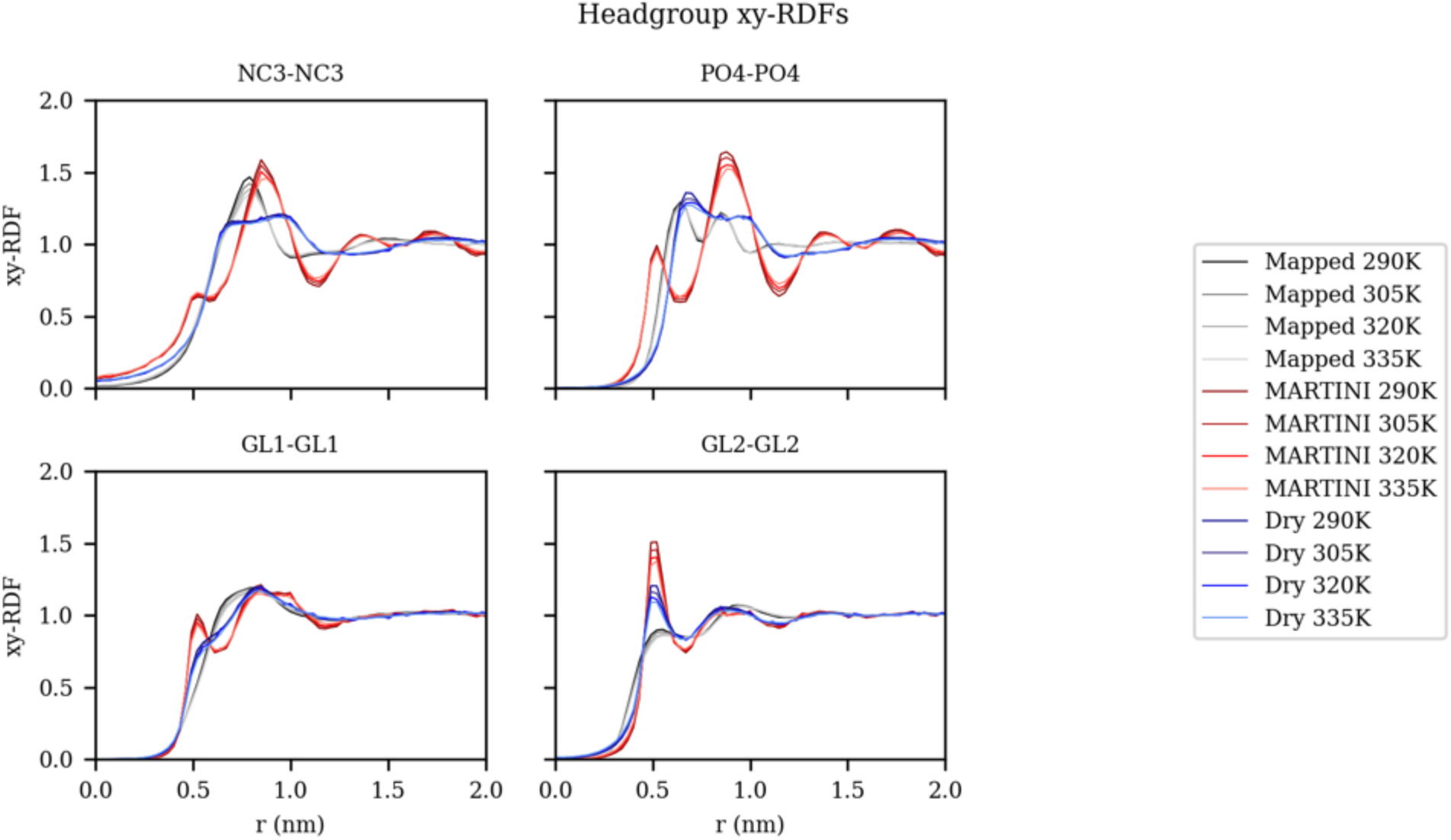
*xy*-projection of the radial distribution function for DOPC head group and glycerol beads comparing the mapped all-atom, MARTINI, and Dry MARTINI models at various temperature averaged per leaflet.

Generally, when comparing the MARTINI RDFs to the mapped atomistic reference for head group beads, it was found that MARTINI 2.0 over-structures and Dry MARTINI under-structures. More specifically, when comparing choline-choline (NC3-NC3) RDFs to the mapped atomistic reference (shown in Fig. 1), MARTINI 2.0 model produced an initial shoulder and secondary features after the first peak, and Dry MARTINI models produced a broader, shorter first peak. Additionally, there was little qualitative variation across the 45K temperature range analyzed, but significant quantitative differences in the first peak heights. The phosphate-phosphate (PO_4_-PO_4_) RDF shows similar behavior: the MARTINI 2.0 model had a significant initial feature before the first peak, and the Dry MARTINI model produced broader peaks. There were significant differences between the DOPC structures.

When considering the DOPC-cholesterol structures, an over-structuring in both the MARTINI 2.0 and Dry MARTINI was found. The glycerol-hydroxyl per leaflet *xy*-projection of the RDF shown in Fig. 2 shows differences between the models and substantial temperature variations in the CG models that are not present to the same extent in the mapped atomistic RDFs.

**Figure 2:**
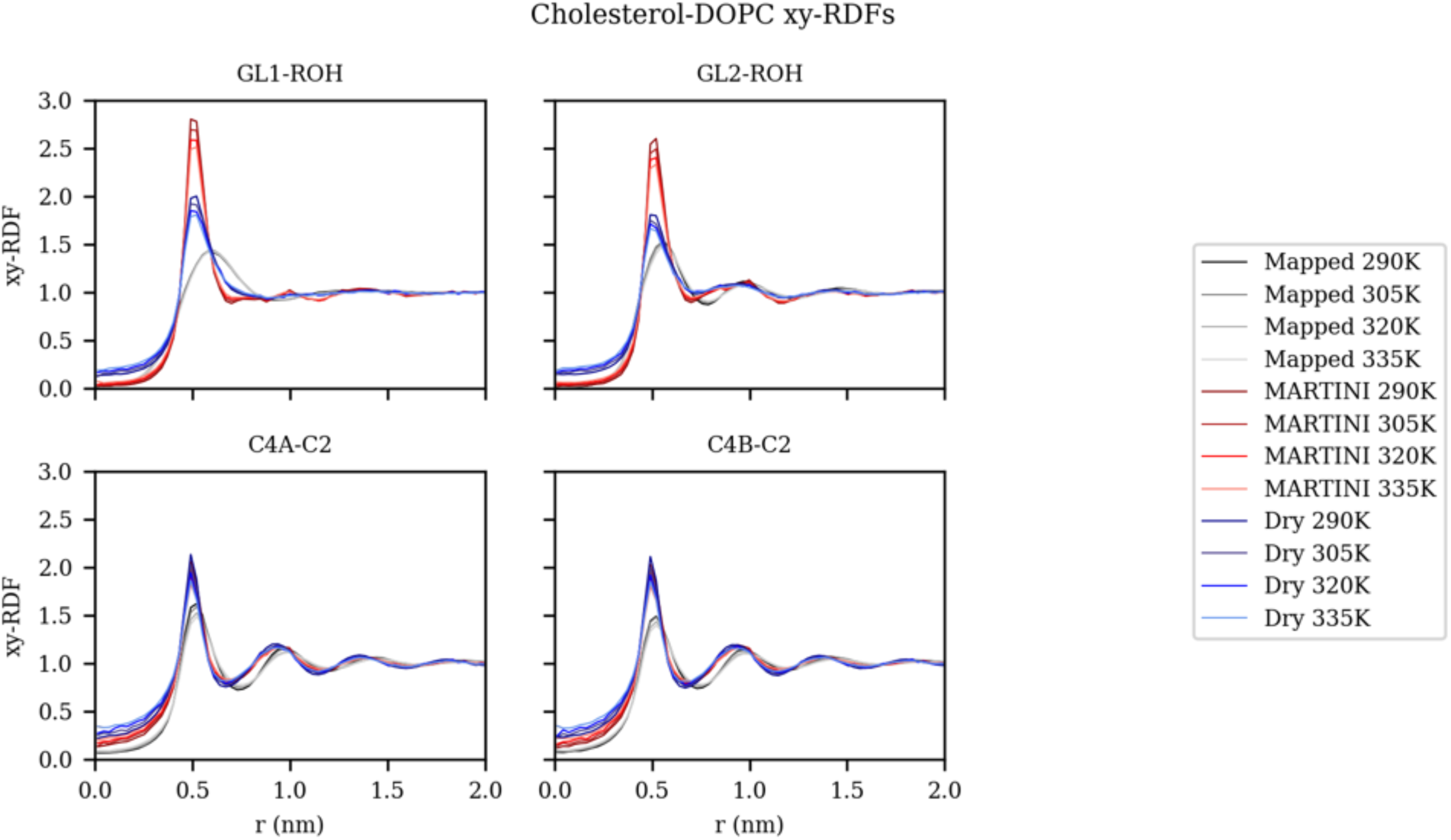
*xy*-projection of the radial distribution function of cholesterol hydroxyl bead (ROH) and DOPC glycerol beads (GL1, GL2) and cholesterol tail (C2) and DOPC terminal carbons (C4A, C4B) comparing the mapped all-atom, MARTINI, and Dry MARTINI models at various temperature averaged per leaflet.

In the remaining RDFs in SI Fig. 1, good agreement between the mapped atomistic reference and both MARTINI models was found. In light of previous analysis of the tail group behavior of MARTINI models, good agreement was to be expected.^34^ Furthermore, the agreement between models suggests that the tail group behavior is largely a packing behavior and one that is well captured by the Lennard-Jones-like interactions of the MARTINI models.

### B. Enthalpy-Entropy Decomposition

By simulating the DOPC/Cholesterol system at various temperatures, a finite difference calculation (see Methods) to decompose the PMF into the enthalpic and entropic contributions was used. Figure 3 shows enthalpy-entropy decomposition of the head group association. The decomposition generally shows there is enthalpy-driven structuring. In MARTINI and Dry MARTINI, it was found there was over-structuring and under-structuring, respectively. The entropic contribution mirrors the enthalpic contribution and provides a driving force to reduce structuring as temperature increases, i.e., the entropic contributions are most negative where the enthalpic contributions are most positive.

**Figure 3:**
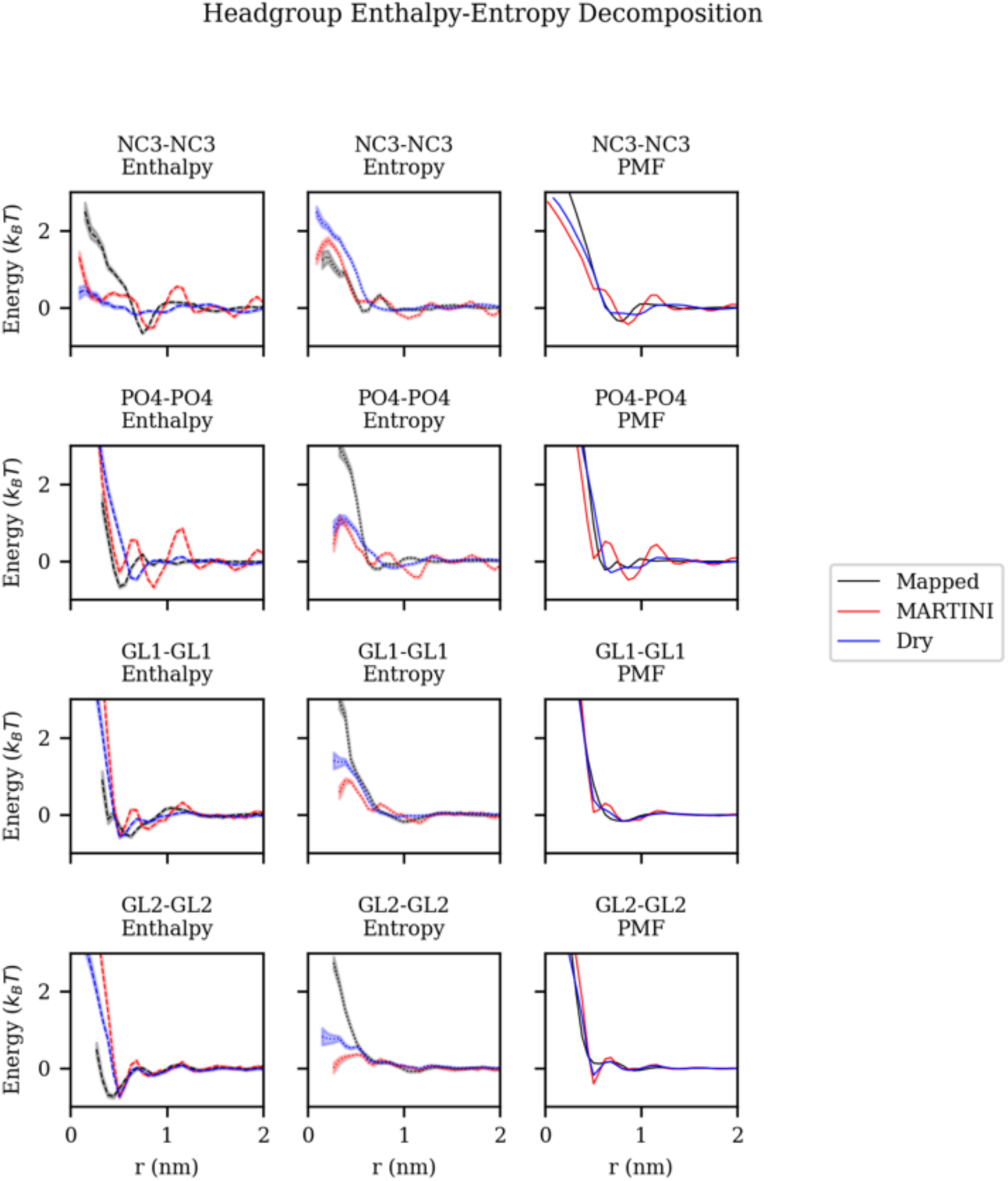
Entropy-enthalpy decomposition of potential of mean force between DOPC head group and glycerol beads comparing mapped all-atom, MARTINI, and Dry MARTINI models.

Based on the DOPC-DOPC headgroup PMFs and corresponding decompositions, the DOPC-cholesterol PMFs were expected to have strong enthalpic contributions that lead to the over-structuring and commensurate entropic contributions to reduce structure as temperature increases. Indeed, enthalpy-entropy decomposition of DOPC-cholesterol interactions shown in Fig. 4 exhibits this behavior. Moreover, the MARTINI 2.0 decomposition has a larger magnitude enthalpy and entropy than the other models at close distances. On the other hand, the MARTINI behavior was the opposite of the calculated behavior of the CG mapped atomistic model. There were small enthalpic contributions to the PMF and significant enthalpic contributions at the less than 0.5 nm distances. The observed behavior suggests that the preferred distance between glycerol beads and hydroxyl bead of cholesterol was entropically driven in the case of the mapped atomistic system and the opposite was true for the MARTINI systems.

**Figure 4:**
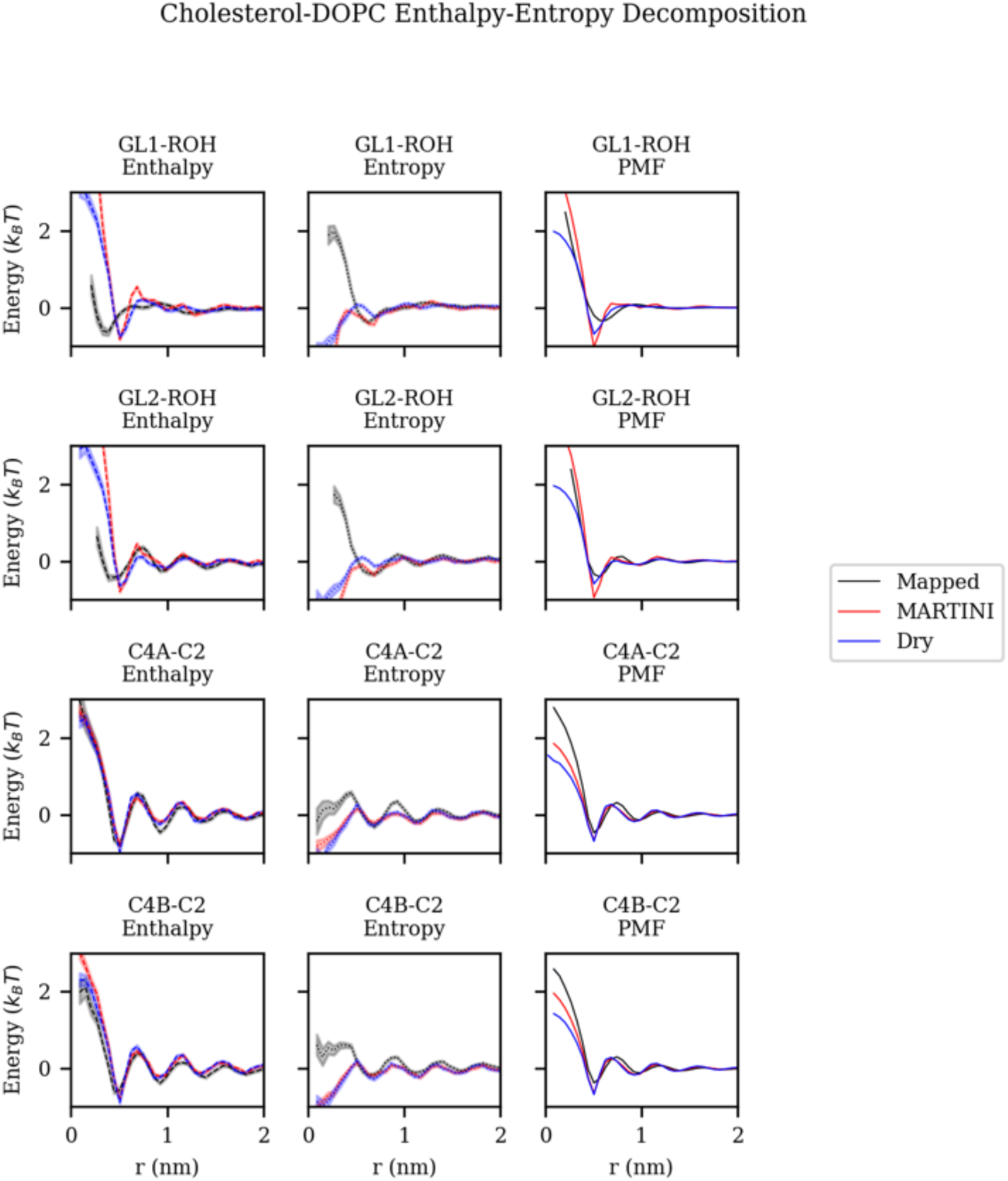
Entropy-enthalpy decomposition of potential of mean force between cholesterol hydroxyl group and DOPC glycerol beads and cholesterol tail group bead and DOPC terminal tail beads comparing mapped all-atom, MARTINI, and Dry MARTINI models.

Finally, as shown in the SI Figures, we again see clear agreement between the mapped atomistic reference and both MARTINI models for the lipid tail CG beads. We expect good agreement between enthalpic contributions to the PMF between the various models because we have generally seen that enthalpy drives the structuring, and there is good agreement between the mapped atomistic, MARTINI, and Dry MARTINI tail RDF structure. Given other studies of the entropy contributed by the tail of MARTINI models, we also expect the entropic contributions to be very similar.^34^

### C. Amphipathic Helix (H0) Lateral Association in the Bilayer

As another test case to assess the differences between atomistic and MARTINI CG models, the lateral association free energy of two embedded helical peptides was estimated by computing the PMF with respect to the CoM distance between the two peptides. The embedded peptides had nonpolar residues pointing toward the hydrophobic core of the bilayer while the polar or charged residues interacted with surrounding solvent or lipid head groups (see Fig. 5A). The system was large enough such that the PMF between the helices decays to 0 kcal/mol at longer ranges. In this configuration, the dipoles of the amphipathic helices were parallel, resulting in the purely repulsive behavior, shown in Fig. 5B. The repulsive decay behavior of the association energy was similar in both atomistic and MARTINI models. However, there was a significant magnitude difference in the repulsion energy between the two models, with the atomistic model reaching repulsive energy of 2 kcal/mol at a CoM distance around 2.0 nm. In contrast, the MARTINI model never came close to that value and, in fact, only one replica reached 2 kcal/mol within the entire region sampled (up to 1.5 nm distance). This result indicates that the MARTINI 2.0 model can greatly increase peptide association behavior in the MARTINI lipid membrane, as revealed by the much too small repulsive behavior in the case of purely repulsive parallel H0 association. Unfortunately, the H0-H0 PMF calculation was too computationally demanding to decompose this behavior into its enthalpic and entropic components. However, one might expect that the spurious behavior of the MARTINI model arises from an incorrect treatment of the decreasing entropy in the PMF as the helices approach one another so that the −*T*Δ*S* term has an incorrect value compare to the CG mapped atomistic result (Δ*S* is negative but considerably too small for MARTINI).

**Figure 5:**
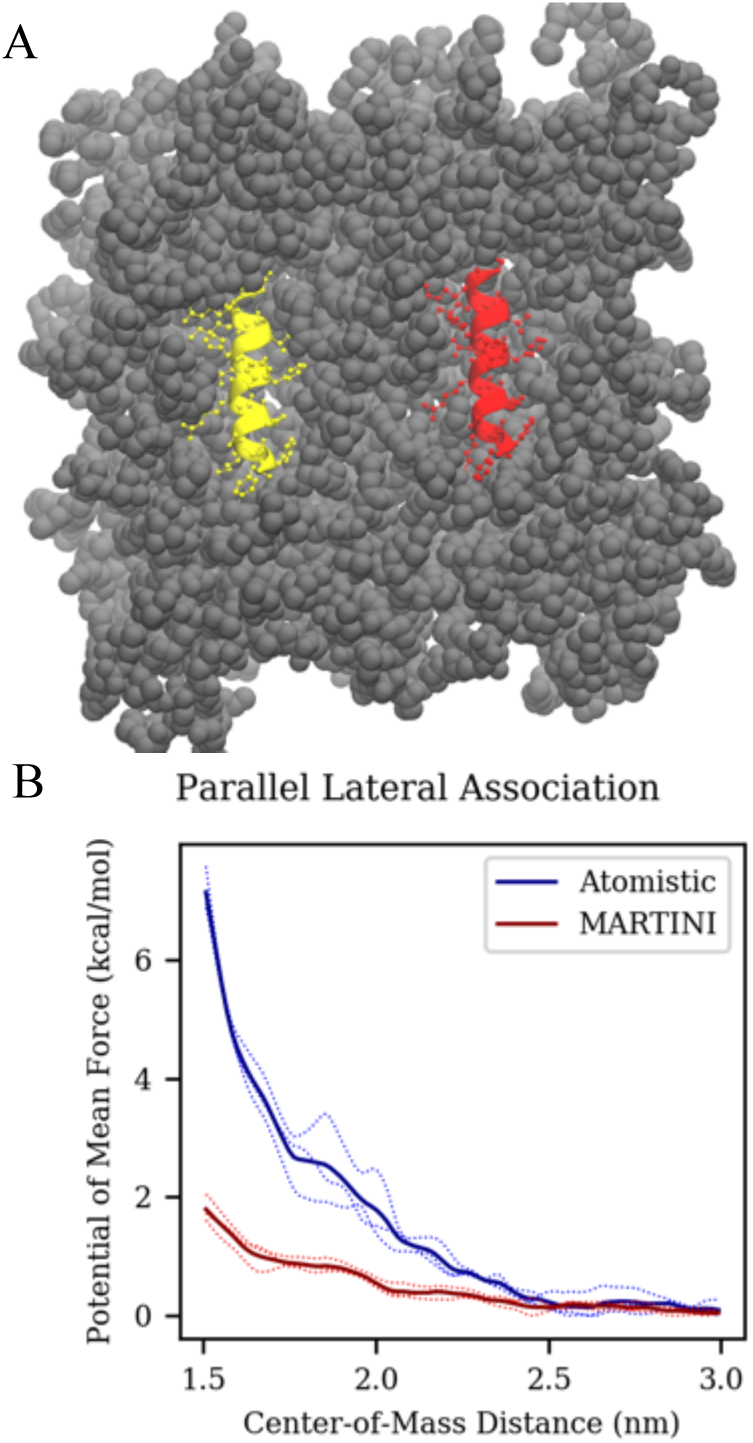
(A) Snapshot of amphipathic H0 helices (red and yellow) embedded in DOPC bilayer (gray) with a center-of-mass distance of 3.0nm, and (B) potential of mean force as a function of center of mass distance between the two embedded helices for the atomistic (blue) and MARTINI 2.0 (red) models (B). Sampling error bars are shown on each curve.

### D. Membrane Undulation Spectrum

Using a 20nm by 20nm lipid bilayer patch of DOPC, the effects of CG mapping and model resolution on the height fluctuation spectrum were investigated. The height fluctuation spectrum is commonly used to access the bending modulus, as described by Canham-Helfrich theory. Additionally, bilayer height fluctuations are also known as an important driving force of, e.g., protein and raft aggregation, because the undulations in the bilayer have the highest number of accessible states when the proteins that dampen fluctuations are closer together. From our calculation of the height fluctuation spectrum (Eqs. 1-3) in Fig. 6, the CG mapping itself did not affect the height fluctuation spectrum (panel A) and, subsequently, did not affect the estimation of the bending modulus at long wavelength.

**Figure 6:**
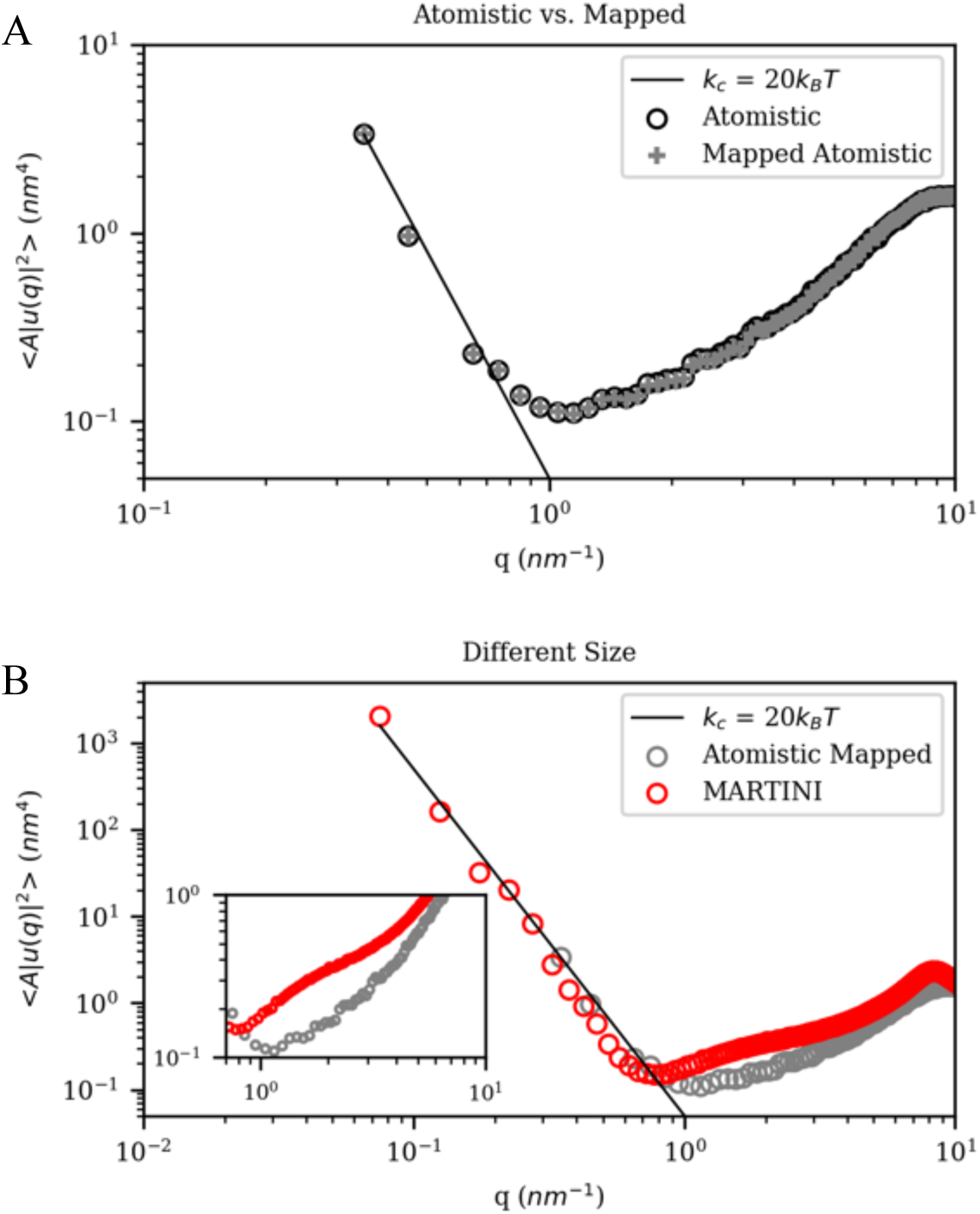
Comparison of undulation spectra. Spectra calculated using the mapped phosphate bead compared to spectrum calculated using phosphorus atom (A). Comparison of spectra calculated from 20nm by 20nm mapped atomistic and 100nm by 100nm MARITNI models (B).

As seen in Fig. 6B, spectra from the CG mapped all-atom and MARTINI 2.0 models qualitatively agreed in the lower *q* (long wavelength) region, which is used to estimate the bending modulus, but they differed somewhat quantitatively in their resulting estimate of bending modulus. The mapped atomistic model had an estimated bending modulus of 22 *k*_*B*_*T* and MARTINI 2.0 bending modulus had a value of 27 *k*_*B*_*T.* MARTINI systems permitted a much larger system (e.g., 100nm x 100nm) to be simulated and showed that the height fluctuation spectrum converges to continuum theory. However, both the shape and the magnitude of the MARTINI spectrum differed very substantially from the mapped atomistic spectrum in the intermediate q regime, i.e., 0.8 nm^−1^ to 6 nm^−1^ (Fig 6B inset and note the logarithmic scale). The intermediate regime lacks a direct correspondence to Canham-Helfrich theory and the assessment of bending modulus, but it is nevertheless on the scale of important membrane-mediated protein interactions. These intermediate scale results suggest that MARTINI simulations of membrane-bound or associated species may be affected by spurious motions in the MARTINI simulations, even when statistically averaged as was done here.

## Discussion

### A. The Driving Forces of Lateral Lipid Association

In Figs. 1-4, we described the lateral structure of a DOPC-cholesterol bilayer and decomposed the enthalpic and entropic contributions to the lateral PMF that gives rise to their association. It was found that there is good agreement in the hydrophobic region of the bilayer but significant disagreement in the head groups in both structure and driving force. This observation is best highlighted by considering the structure and driving force between glycerol beads of DOPC and the hydroxyl-group bead of cholesterol. Figure 4 showed that not only are both MARTINI systems significantly over-structured, but also that the interaction was enthalpically driven in the MARTINI systems but entropically driven in the CG mapped atomistic system. Thus, in addition to being driven by physically incorrect features of the interactions, the temperature dependence of the MARTINI systems will also be inherently wrong due to an incorrect treatment of entropic effects.

### B. Lateral Association of Embedded Amphipathic Helices

Amphipathic helices are an important motif for membrane protein targeting and assembly.^35-36^ The association energy between two amphipathic helices is an essential test case because it is likely affected by the over-structuring behavior seen in Fig. 1 and 2 and the membrane height fluctuation differences seen in Fig. 6. More specifically, the larger undulations in the MARTINI model could cause a greater effective attraction between embedded helices as the closer association of helices cause a decreased dampening of membrane fluctuations.^37^ Additionally, it may also be affected by the solvent behavior of the 4 waters to 1 bead MARTINI mapping as solvating the amphipathic helices is different in the two models as well. Finally, it is a multi-component system and suffers from the lateral association issues seen in Fig. 4 (i.e., the model may be erroneously attributing the repulsion to either enthalpic or entropic effects). Therefore, as noted earlier, Fig. 5 showed the MARTINI repulsive free energy between the two amphipathic helices to be significantly underestimated. This result, and those of Figs 1-4, should also be noted in light of ambitions to simulate via MARTINI very complex membrane-protein systems in the coming years.^38^

### C. Membrane Undulation Spectrum

It was found that the CG mapping has little or no effect on the height fluctuation spectrum and the subsequent estimate of bending modulus. We attribute this to our original definition of the bilayer mid-plane, which was the average position of the phosphorus atoms in each leaflet. Thus, when the phosphorus atom is mapped with the bonded oxygen atoms to the phosphate bead, there was relatively little change in the estimation of the midplane, and thus there was little to no difference due to representability. This would not be true if our definition of the midplane was more sensitive to CG mapping. For example, if the tail carbon was chosen, there would be a significant effect because the position of the last tail carbon is significantly more dynamic than the last tail bead. As a result, the mapped spectrum would have an effective filter upon the short wavelength (high q) fluctuations, which should not affect the estimate of the bending modulus. However, this should only act as a short wavelength filter, and the undulation spectrum should converge to *q*^−4^ behavior as expected from Canham-Helfrich theory.

However, the role of model resolution on the height fluctuation spectrum is of importance when considering the growing number of CG lipid models. Figure 6 showed that the height fluctuation spectrum at long wavelengths (low *q*) does not have significant effects due to model resolution and does converge to *q*^−4^ as expected from continuum theory. When considering the role of CG mappings and the definition of the bilayer midplane, this result was perhaps not surprising. However, a surprising finding here was the behavior in the intermediate *q* regime of the height fluctuation spectra. The intermediate regime corresponds to distances on the order of membrane-mediated protein-protein interactions where the role of lipid bilayer fluctuations is exceedingly important (see, e.g., ref ^39^). There were much higher average height fluctuations in the case of the MARTINI 2.0 model spectrum with a qualitative shape difference as compared to the CG mapped atomistic spectrum. This suggests that the physical origin of these fluctuations may not even be the same. Indeed, we expect the solvent and the interactions between adjacent lipids to play a significant role in the nature of intermediate length scale lipid height fluctuations. When considering the case of the MARTINI 2.0 model, these fluctuations are not as dampened by the solvent and interactions between adjacent lipids. While the reduced representation of water in the MARTINI 2.0 model is not a primary focus in this work, this primitive representation of the liquid and its solvation of the membrane cannot be ignored when considering the membrane fluctuations and potentially over-stabilizing (or not adequately damping) lipids moving outward from the membrane. It may be that an overly attractive force between MARTINI water and lipids pulls lipids out of the membrane and stabilizes local fluctuations. but this and related questions can be a focus of future research.

## Conclusions

The MARTINI model, as an example of a largely “top-down” CG approach, has become widely adopted due to its ease of use and apparent applicability to a wide variety of systems. In that context, the lateral ordering and driving forces of a relatively simple case of DOPC and cholesterol bilayer were investigated by calculating the per leaflet *xy*-projection of the RDF and the enthalpy-entropy decomposition of the corresponding PMF. It was found that the hydrophobic region of the bilayer is described almost as well by the MARTINI models as a CG mapped atomistic system. However, there was significant disagreement in the head group region, both in structuring, and the enthalpy-entropy decomposition. The cross interactions between the components were erroneously driven by enthalpy as opposed to entropy, as seen in the CG mapped atomistic system. This result may give pause to a growing community of modelers investigating increasingly complex systems at the CG scale with such top-down approaches, often across a wide composition range and a varying temperature scale. The enthalpy-entropy balance is typically delicate in biomolecular systems. The misattribution of enthalpy-entropy in the interactions may give rise to erroneous conclusions when the CG model is used to simulate systems at different state points, or even those state points for which it was initially parameterized in a top-down “fitting” approach. As shown in this work, such problems may arise even in something as basic as helix-helix association free energy in a membrane

Additionally, the role of CG mapping the underlying model on the membrane fluctuations was investigated. At low wavenumbers (or long wavelengths), the spectrum is only quantitatively affected by mapping or model resolution, while at the intermediate wavelengths of 0.8 nm^−1^ to 6 nm^−1^, there were large deviations between MARTINI 2.0 and the CG mapped atomistic model. These differences were not due to the representability of the height fluctuation spectrum, but we suggest that these differences are more likely due to model differences in the inherent stability of undulating membranes. The solvent also plays a significant role in lipid bilayer stability. Indeed, the four waters to one CG bead mapping in the case of the MARTINI 2.0 does not appear to provide the correct energetic barriers to bilayer undulations, resulting in significantly higher magnitude fluctuations in the MARTINI model. Such erroneous membrane fluctuations would, among other things, directly impact membrane-mediated interactions of associating proteins. Indeed, as noted earlier when we investigated the lateral association of two embedded amphipathic helices we found an underestimation of their repulsion energy.

By making a top-down CG model such as MARTINI more in tune with the quantities and insights of statistical mechanics, it may be possible to expand the range in which such models can be trusted to realistically model complex biomolecular and other soft matter systems. The steps taken in this work to explore these features may also hopefully lead to additional efforts to parameterize such models to bring them into greater agreement with the known laws of physics and, in particular, those of statistical mechanics.

## ACKNOWLEDGMENTS

This research was supported by the National Institute of General Medical Sciences (NIGMS) of the United States National Institutes of Health (NIH) under NIH award number R01-GM063796. The computations were supported by the University of Chicago Research Computing Center (RCC).

## TABLE OF CONTENTS GRAPHIC

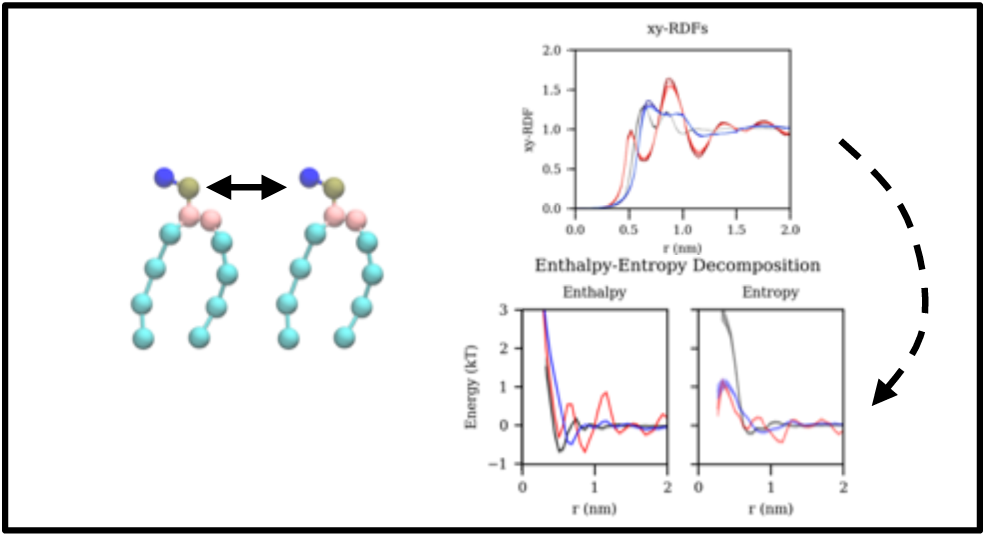

## Notes

### Competing Interest Statement

The authors have declared no competing interest.

