## Supplemental File for "Coarse-grained Force Fields from the Perspective of Statistical Mechanics: Better Understanding the Origins of a MARTINI Hangover"

It has been discussed elsewhere that the entropy of the lipid tail groups is well captured by the MARTINI model.[1] As such, we expect the radial distribution functions (RDFs) and the atomistic temperature behavior to be recapitulated. The following figures presenting the tail CG bead RDFs and corresponding enthalpy-entropy decompositions, which are calculated as described in the Methods section of the main document.

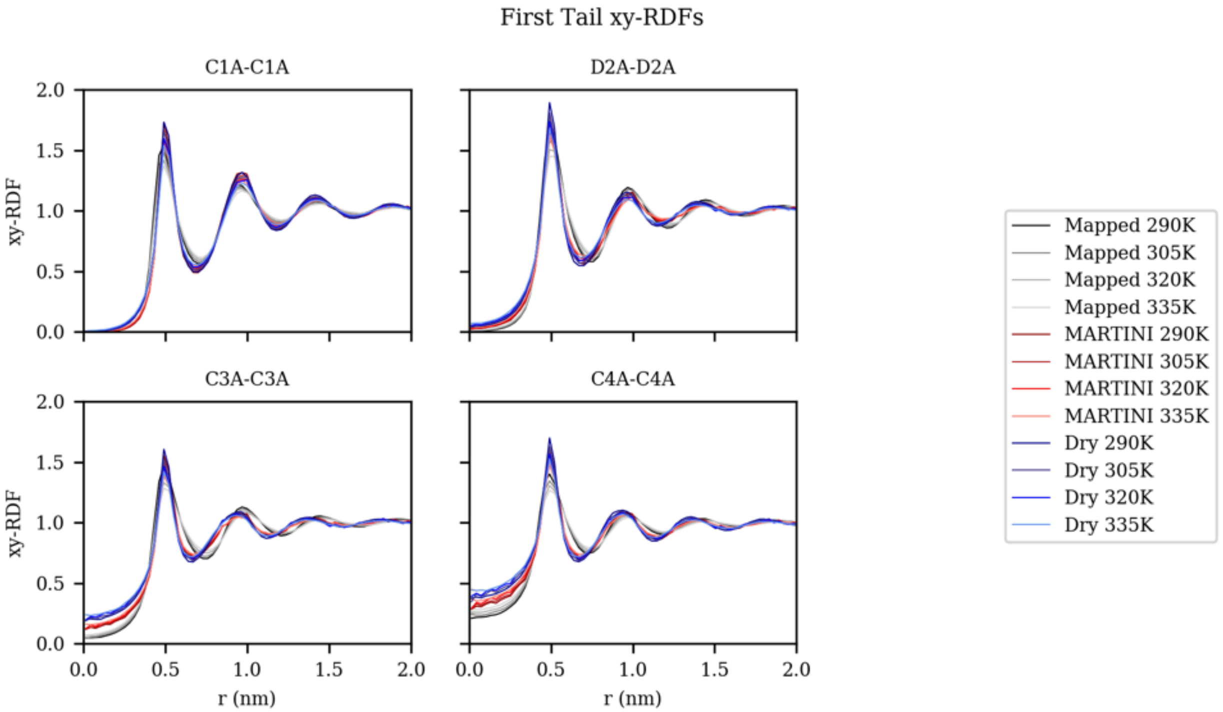


**SI Figure 1:** *xy*-projection of the radial distribution function for DOPC first tail CG beads comparing the CG mapped all-atom, MARTINI, and Dry MARTINI models at various temperature averaged per leaflet.

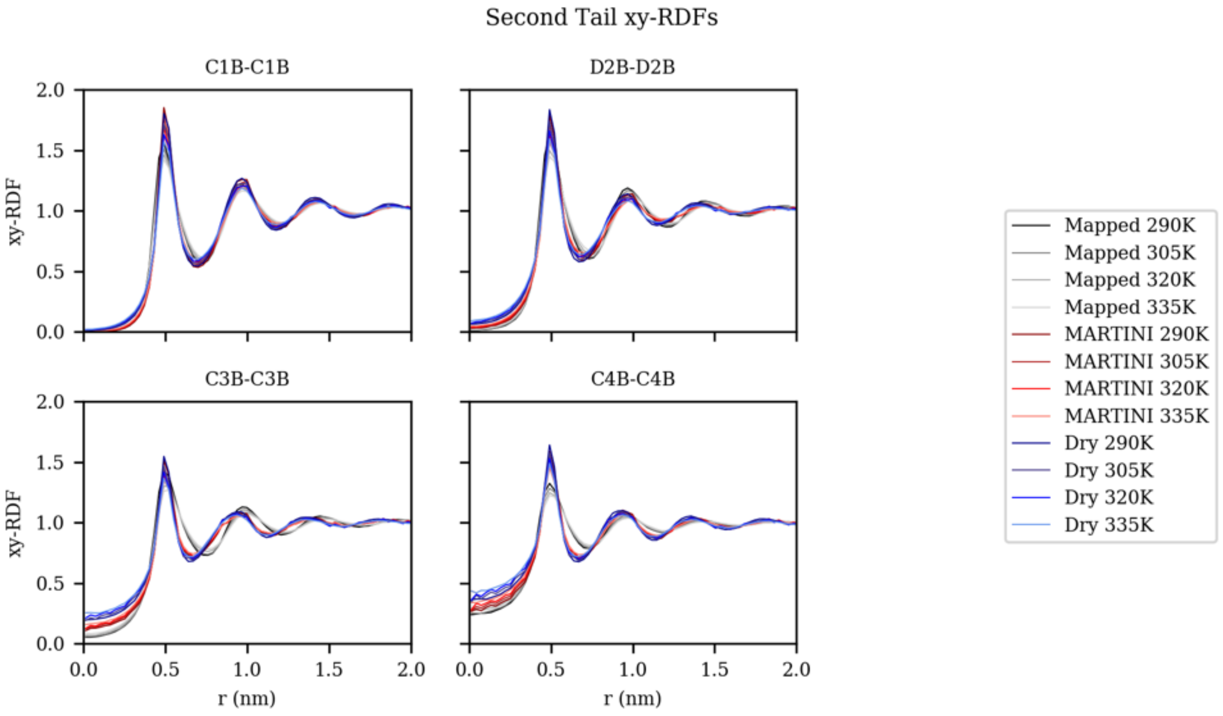


**SI Figure 2:** *xy*-projection of the radial distribution function for DOPC second tail CG beads comparing the CG mapped all-atom, MARTINI, and Dry MARTINI models at various temperature averaged per leaflet.

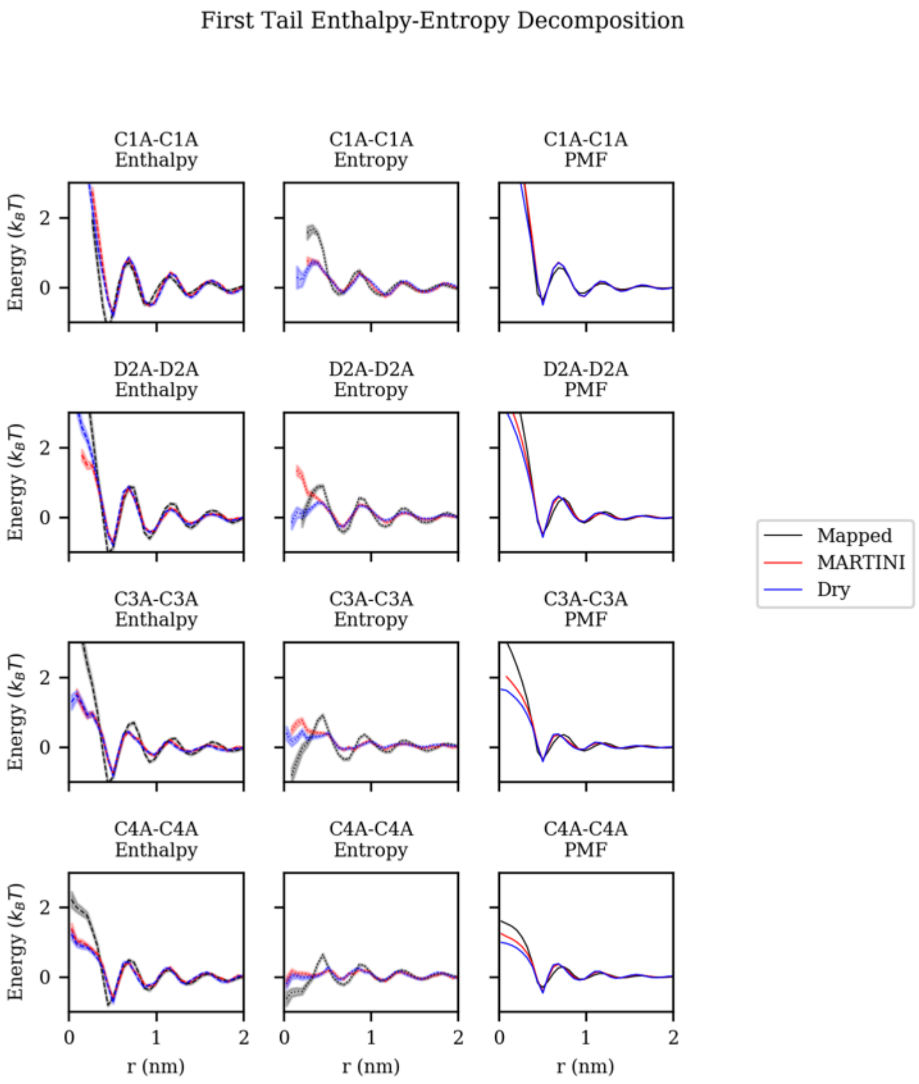


**SI Figure 3:** Entropy-enthalpy decomposition of potential of mean force between first tail CG beads of DOPC comparing CG mapped all-atom, MARTINI, and Dry MARTINI models.

Second Tail Enthalpy-Entropy Decomposition

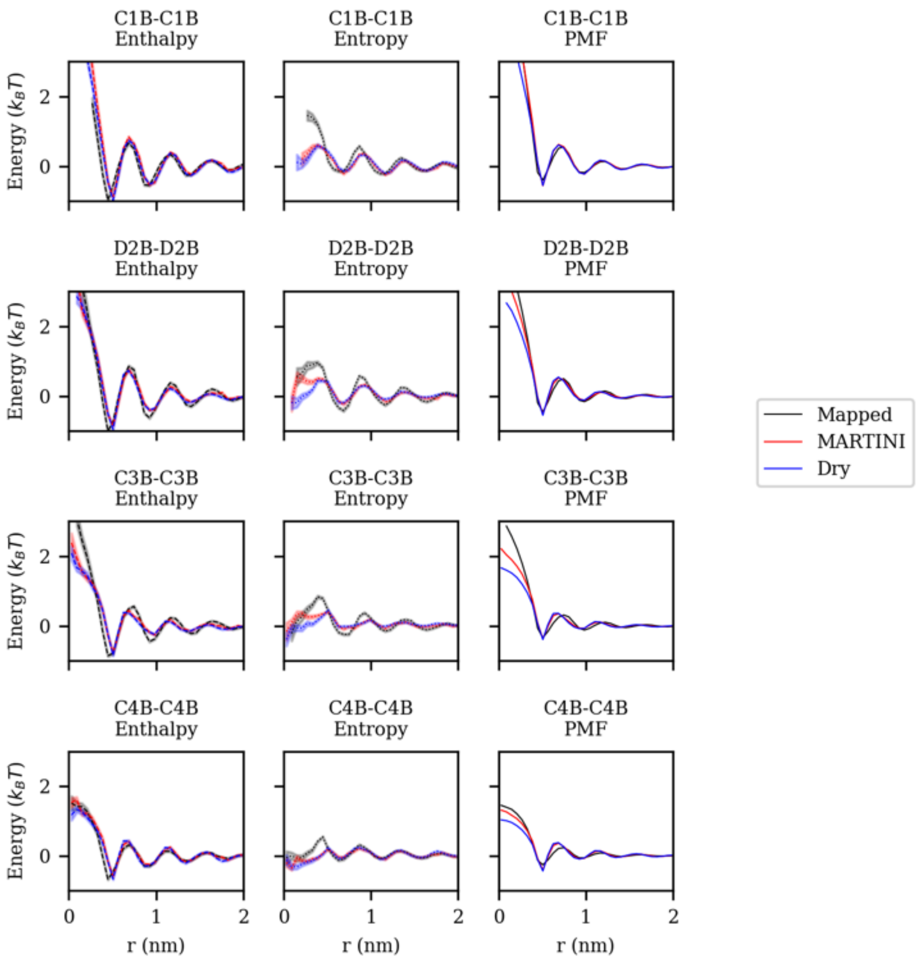


**SI Figure 4:** Entropy-enthalpy decomposition of potential of mean force between second tail CG beads of DOPC comparing CG mapped all-atom, MARTINI, and Dry MARTINI models.
